# Triplex DNA and inverted repeats cause long-read sequencing bias against simple satellite DNA

**DOI:** 10.64898/2026.07.13.738322

**Authors:** A. Bernardo Carvalho, Bernard Y. Kim, Fabiana Uno

## Abstract

We recently showed that Oxford Nanopore Technologies (ONT) and Pacific Biosciences (PacBio) have very strong sequencing bias against simple satellites, probably caused by single-stranded DNA folding into non-canonical (non-B) structures during sequencing. Here we extend these observations by computational and experimental approaches in the *Drosophila* and human genomes. We found that (i) only a small subset of simple satellites cause sequencing bias; many satellites (*e.g.*, (ACTGGG)n) are benign and easily sequenced. (ii) The biases most likely are caused by two distinct non-B DNA structures: triplex DNA formed by some, but not all, AG-rich satellites (only those predicted to form strong mirror repeats), and hairpins formed by some, but not all, AT-rich satellites (only those predicted to form fairly strong inverted repeats). (iii) The correlation between the predicted stability of these non-B structures, and the strength of sequencing bias indicates that non-B DNA is indeed the culprit. (iv) The likely source of these non-B structures is single-stranded DNA formed during ONT and PacBio sequencing, and hence its removal might solve the bias. We tested this by adding single-strand binding protein to ONT sequencing, and found that it irreversibly kills the flow cells. (v) A recent sequencing effort in *Drosophila melanogaster* using very high depth Ultra-Long ONT sequencing (967×) still failed to assemble many genes located near satellite blocks. Brute-force will not solve the problem; instead, further investment is needed by the sequencing companies to achieve truly unbiased sequencing.

## Introduction

Long-read sequencing technologies — Oxford Nanopore Technologies (ONT) and Pacific Biosciences (PacBio) — are widely regarded as nearly free of sequence-composition bias, which alongside their read lengths, underlies their success in producing telomere-to-telomere (T2T) assemblies. However, Carvalho et al. (2026) recently found that both technologies systematically fail to sequence and assemble exons of *Drosophila melanogaster* Y-linked genes that lie near simple satellite blocks. This bias was noticed by Carvalho et al. (2016) while analyzing the first *Drosophila* long-read dataset (Kim et al. 2014), and at that time was incorrectly attributed to a DNA purification protocol. Later on, Nurk et al. (2022) noticed that small blocks of GA-rich sequences (*e.g.*, a 256 bp (AAAGG)n block in the human chromosome 8) caused assembly breaks due to strongly reduced coverage. The bias is also very strong in *Drosophila*, being able to cause “missing exons” even at high coverage (200 ×). Carvalho et al. (2026) proposed that the underlying cause of both cases is non-B DNA, specifically, triplex DNA (H-DNA) formed when single-stranded DNA, generated during sequencing, folds back onto the duplex near the sequencing site. The main evidence was that *Drosophila* Y-linked low-coverage exons, but not normal-coverage ones, are highly enriched in mirror repeats, a requirement for triplex formation ((Mirkin et al. 1987; Hisey et al. 2024). Other non-B DNA motifs, such as quadruplex DNA and Z-DNA, do not show this regular association.

That study opened several questions, which we aim to answer here using computational and experimental approaches in the *Drosophila* and human genomes. First, Carvalho et al. (2026) examined only low-coverage regions, and hence could not tell whether all simple satellites, or only a subset, disrupt sequencing. Second, while triplex DNA is a compelling explanation for AG-rich-satellites, it cannot account for low-coverage regions such as *kl-5* exon 14, which contains only (AATATAT)n; for the sake of simplicity we will hereafter call the satellites by their monomers, *e.g.*, AATATAT. This satellite is a perfect mirror repeat, but triplex formation additionally requires homopurine / homopyrimidine strands (Mirkin et al. 1987), which AT-only satellites lack. This implies that a second, distinct mechanism must be at work. Third, if non-B structures indeed arise from single-stranded intermediates during sequencing, then removing that single-stranded DNA should mitigate the bias; this offers both a direct test of the hypothesis and a potential route for a solution. Finally, it remains unclear whether the problem could simply be out-sequenced: a recent ultra-deep effort in *D. melanogaster* (Liu et al. 2025; 967x ultra-long ONT plus 107x HiFi) claimed to have achieved a near-T2T Canton-S assembly, raising the possibility that brute-force coverage might suffice.

## Results

### Do all simple satellites cause sequencing bias?

We approached this question by searching for all simple satellite blocks in the *D. melanogaster* Release 6 (“R6”) assembly (Hoskins et al. 2015; Ozturk-Colak et al. 2024), and measuring PacBio HiFi (hereafter, “HiFi”) read coverage across these regions. We focused on the gene regions annotated in R6 (Methods) because they are less repetitive than the whole genome and therefore less prone to mapping problems. We used HiFi reads because they are more sensitive to satellites (Nurk et al. 2022; Carvalho et al. 2026; Supplemental Fig. S1). Following Carvalho et al. (2026), we searched for perfect satellite repeats using the script *find_tandem_repeats_v2.py*. To avoid confounding the analyses with satellite blocks that are too small, we required at least 250 bp of perfect satellite repeats. We adopted this cut-off because the 256 bp AAAGG sequence caused an assembly break in the human genome (Nurk et al. 2022). We identified 32 gene regions containing satellite blocks larger than 250 bp (Supplemental Table S1). This list includes many low-coverage genes - and the corresponding satellite sequences - identified by Carvalho et al. (2026), such as *kl-3* / AATATAT, *Ppr-Y* / AAGAG, and *CG42402* / AAATATAT. This list also includes several other genes and satellites not known to be associated with low read coverage, such as *Cda4* / ACTGGG. In many of the latter cases, the R6 gene sequence is continuous (*i.e.*, not interrupted by the NNN blocks used to mark assembly gaps). These two observations suggest that several simple satellites do not cause sequencing problems, although it should be noted that R6 was produced from Sanger reads rather than long reads. Indeed, as shown in Fig. 1B and Fig. 1C, when we aligned HiFi reads to these gene regions, the coverage is uniform and close to the expected value. Although in some cases read mapping and other technical issues prevented an unambiguous answer, several simple satellites are clearly benign. For example, the ACTGGG 1968 bp block in *Cda4* and the AAATAAT 917 bp block in *Osi19* did not cause any coverage drop (Fig. 1). So it is clear that the ability to disrupt long-read sequencing of AAAGG is not shared by all simple satellites. For the sake of simplicity we will refer to satellites like AAAGG as “toxic” and those like ACTGGG as “benign”. Interestingly, the toxic *kl-3* / AATATAT and the benign *Osi19* / AAATAAT satellites are very similar (both are 7 bp, 100% AT), yet have different effects in long-read sequencing. This indicates that base composition *per se* is not the explanation of the sequencing bias.

**Figure 1.**
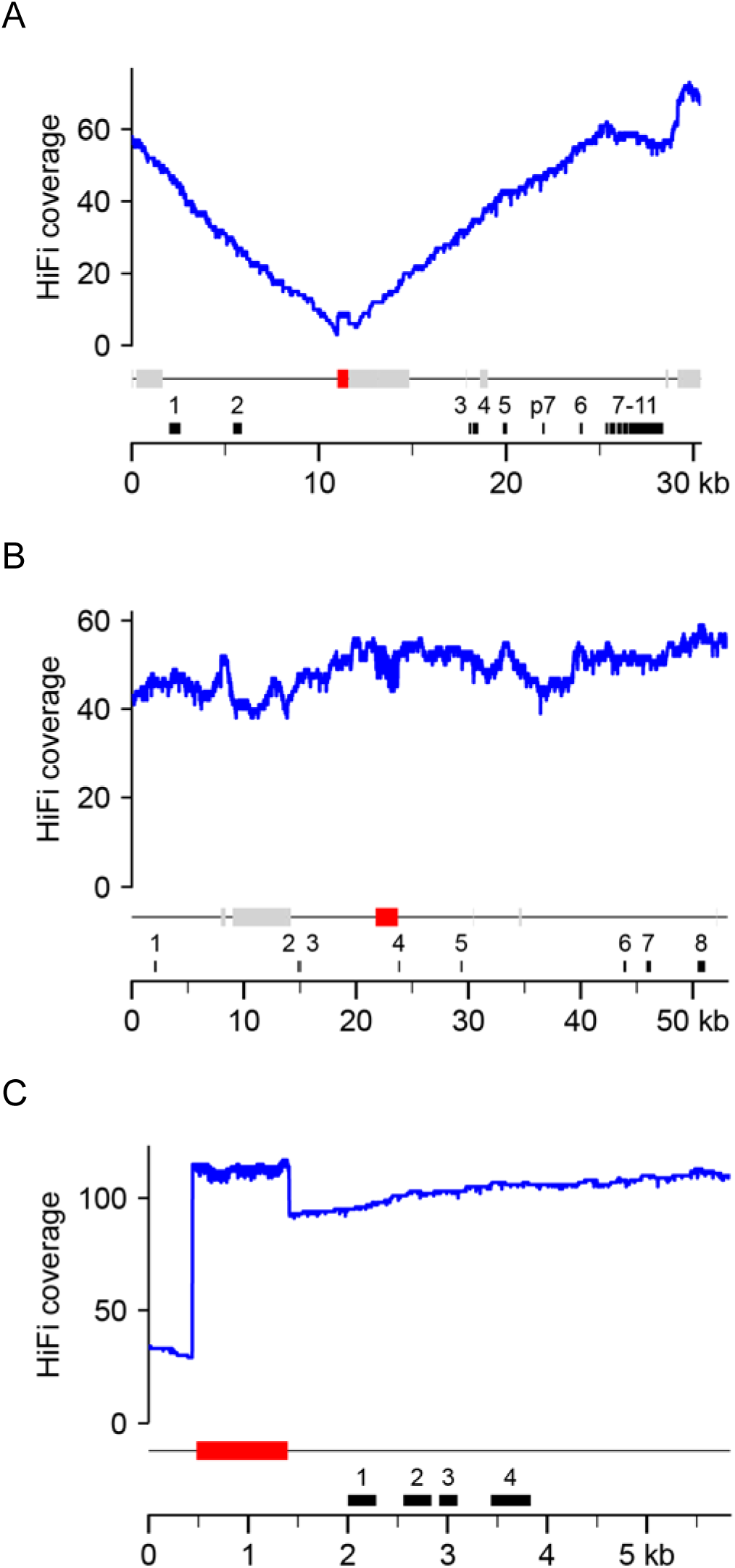
Toxic and benign satellites in R6 gene regions. HiFi coverage profiles of the *CG42402* (A), *Cda4* (B) and *Osi19* (C) gene regions are shown. Exons of the genes are in black (“p7” stands for a pseudogene of exon 7), transposable elements are in light grey, and the main satellites in red. A) *CG42402* region, containing a 576 bp block of the satellite AAATATAT. R6 collapsed it, its actual size being ∼50 kb (Fig. 2B). *CG42402* is autosomal, and HiFi coverage is 97x; AAATATAT is clearly toxic. B) *Cda4* region, containing a 1968 bp block of the satellite ACTGGG. The *Cda4* gene is X-linked, so we would expect ∼ 50x coverage. Note that the satellite does not cause any coverage drop. C) *Osi19* region, containing a 917 bp block of the satellite AAATAAT. The *Osi19* gene is autosomal, so we would expect ∼ 100x coverage. Note that the coverage is essentially flat, except for the increase in coverage in the satellite region, resulting from assembly collapse in R6 (Fig. 2C shows that the actual block is ∼30 kb long). ACTGGG and AAATAAT are benign.

Not all satellites listed in Supplemental Table S1 as not causing coverage drops are benign: as detailed later, while ∼250 bp of some toxic satellites (*e.g.*, AAGAG) completely disrupt HiFi sequencing, others (*e.g.*, AAATATAT) only cause major problems with blocks larger than 1-2 kb (Supplemental Fig. S2).

### A genome-wide search for toxic and benign satellites

The approach of the previous section would miss satellites that are not closely associated with genes. Furthermore, it relies on the R6 assembly, a patchwork of decades of sequencing and finishing efforts that employed a variety of methods, which raises the possibility of bias. More importantly, R6 collapsed several large satellite blocks (*e.g.*, Fig. 1A vs. Fig. 2B). Hence, we extended that approach by searching for simple satellites in assemblies produced with ONT 10.4, PacBio HiFi, and PacBio LILAP reads (Jia et al. 2024; Kim et al. 2024; Shukla et al. 2025), all assembled with *hifiasm* (Cheng et al. 2026). The LILAP protocol replaces T4 ligase with the Tn5 transposase in the main step of adaptor-target DNA ligation; Carvalho et al. (2026) found that *Drosophila* LILAP reads are less biased, probably due to their smaller insert size (∼5 kb). The three read datasets came from the same reference strain of *D. melanogaster* (iso-1) used in the R6 assembly. Among the four assemblies, HiFi-hifiasm had the largest total amount of simple satellites (3.1 Mbp), followed by the ONT-hifiasm (2.7 Mbp; Supplemental Table S2). However, ONT-hifiasm had by far the best representation of toxic satellites (Supplemental Fig. S3) and, unless otherwise noted, was used in all analyses.

**Figure 2.**
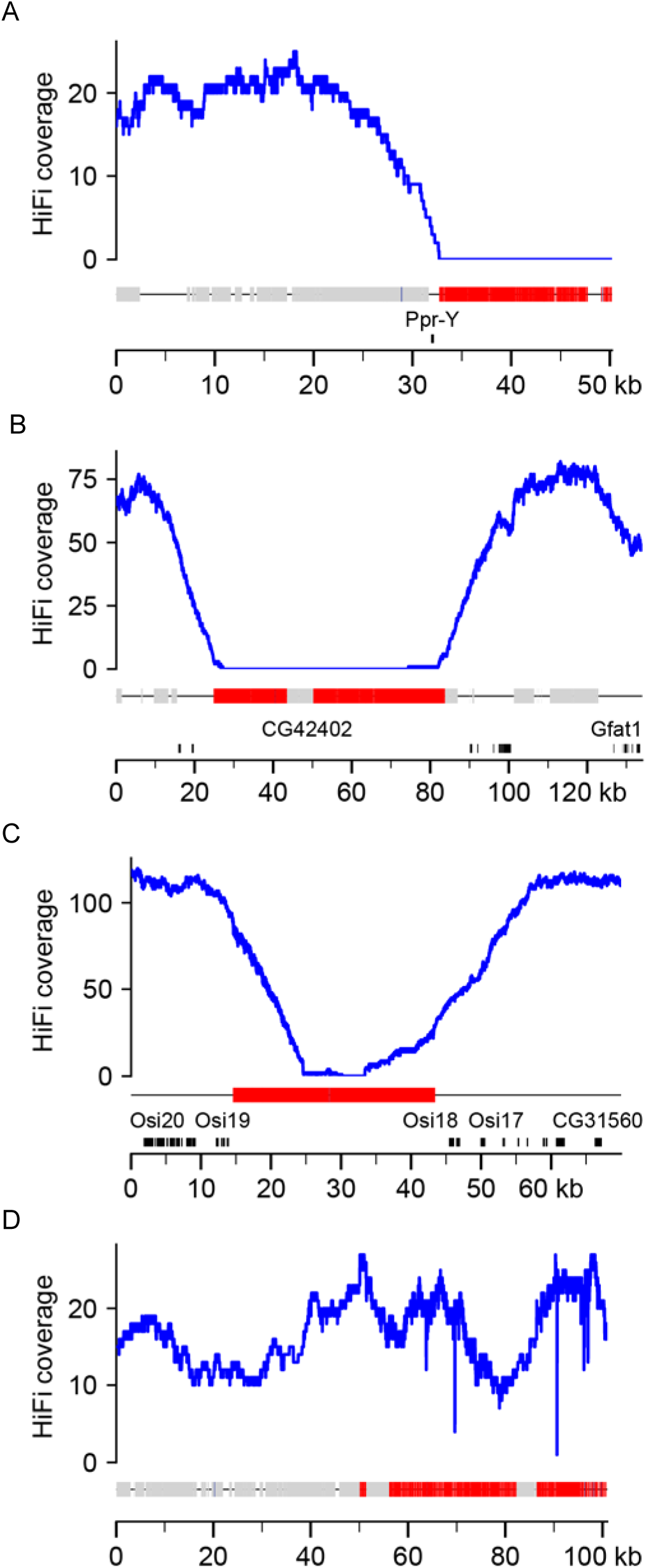
Coverage profile patterns of toxic and benign satellites (ONT assembly). A) coverage profile of AAGAG (a very toxic satellite): HiFi coverage drops to zero before entering the satellite block. B) AAATATAT, another toxic satellite; note that HiFi coverage quickly drops to zero inside the satellite block. Panels C and D show two benign satellites, AAATAAT and AAAT: HiFi read coverage penetrate 10kb-15kb inside the satellite block, and may not even reach zero. The key parameter that distinguishes toxic and benign satellites is how much the HiFi coverage penetrates the satellite block (Supplemental Methods; Supplemental Fig. S5). Gene exons are labelled in black, transposable elements in light gray, and the main satellite in red.

As expected, we got a large list of simple satellite regions. We found 2,648 perfect satellite blocks (if we merge those located within 100 bp of each other, there are 1,996 blocks), belonging to 65 different satellites (Supplemental Table S3). Many of the 65 satellites are very rare and occur in small blocks (*e.g.*, 18 of them occur once or twice, and have a total size of 11 kb), but abundance in the assembled genome is a poor indicator of true genomic abundance, due to sequencing bias and collapse during assembly. We initially computed aggregate statistics for each satellite (*e.g.*, median read coverage) to identify low-coverage satellites, but found this too error-prone: several satellites co-occur (*e.g.*, Supplemental Fig. S3 bottom), and in these cases the detrimental effect on read coverage from one satellite “spreads” to its neighbors. Instead, we examined the HiFi read coverage profile of individual satellite blocks in detail, in order to identify benign and toxic satellites. This block-by-block approach allowed us to control for co-occurrence, collapsed regions, and other sources of artifacts (blocks with significant amounts of co-occurring satellites were excluded from the analysis). Given the large number of satellite blocks (2,648), and the fact that all toxic satellites identified by Carvalho et al. (2026) are either 100% AG or 100% AT, we focused on them. Below we present the general findings for both AT-only and AG-only satellites, and in the next sections we detail them separately.

We found that toxic satellites cause a coverage drop ∼15 kb before the start of the satellite block, presumably because DNA fragments containing even a small amount of these satellites cause sequencing failure (Fig. 2). The HiFi coverage gradually declines as one approaches the start of the satellite block, reaching zero in its vicinity (AG-only satellites; Fig. 2A), or at most a few kb inside it (AT-only satellites; Fig. 2B). The case of benign satellites (Fig. 2C and Fig. 2D) is a bit more complex due to mapping issues. Namely, reads originating entirely within a long perfect repeat cannot be uniquely mapped, and hence even a completely benign satellite can show reduced coverage if its block size approaches or exceeds the HiFi read size (∼16 kb; Shukla et al. 2025). This effect is hard to predict because satellite blocks vary in heterogeneity, with internal variants serving as “anchors” that allow read mapping to penetrate into the block. When such drops occur in benign satellites, they start closer to the satellite block, and HiFi coverage typically penetrates at least ∼15 kb into the satellite block (Fig. 2C).

The above conclusions about toxic and benign satellites are independently supported by examining the satellite content of raw reads, which circumvents the effects of read mapping and assembly collapse, as follows. We obtained the total amount of each satellite in HiFi and Illumina reads (the latter assumed to represent the genome ground truth) and calculated their ratio (Supplemental Results). We would expect that for toxic satellites the Illumina / HiFi ratio would be very high, whereas for benign satellites it would be closer to 1. This was indeed the case (Table 1; Supplemental Table S4). There are several caveats here (Supplemental Results), in particular the use of Illumina as the ground truth, since Illumina suffers from bias against AT-rich and GC-rich regions (Oyola et al. 2012; Wei et al. 2018; see also the Supplemental Results section “Detecting satellite toxicity in raw reads”). Note, however, that the three satellites shown in Table 1 have a similar, neutral GC composition (40%-50%). Overall, the difference between benign and toxic satellites shown in Table 1 is so extreme that it can hardly be attributed to anything other than the HiFi sequencing bias against toxic satellites.

**Table 1.** Satellite toxicity in raw reads. The values are the total satellite amount.

| Satellite | Type <sup>a</sup> | Illumina bp <sup>b</sup> | HiFi bp | Illumina / HiFi ratio |
| --- | --- | --- | --- | --- |
| AAGAG | toxic | 13,690,900 | 3,090 | 4431 |
| AAGAGAG | toxic | 1,701,917 | 0 | ∞ |
| AAGGAG | benign | 821,052 | 167,586 | 5 |

<sup>a</sup> The type of satellite (benign vs. toxic) was independently inferred from read coverage profiles (Fig. 2 and Fig. S4)
<sup>b</sup> Illumina reads downsampled to 100 × (the same depth of the HiFi reads).

To summarize, coverage drops caused by toxic satellites are directly caused by negative sequencing bias, whereas those from benign satellites are mostly due to mapping issues.

### Triplex-DNA formation, and not base composition, causes sequencing bias of AG-rich satellites

Three AG-only satellites yield reliable coverage profiles (*i.e.*, have at least one large block in the ONT- hifiasm assembly in which they do not co-occur significantly with other satellites): AAGAG, AAGAGAG, and AAGGAG. The first, AAGAG, is one of the most abundant *D. melanogaster* satellites, accounting for ∼11 Mbp of the male haploid genome (Lohe et al. 1993); only 178 kb were assembled with ONT reads (Table 2). Its coverage profile (Fig. 2A; Supplemental Supplemental Fig. S4) shows that it is invariably very toxic: HiFi read coverages begin to fall ∼15 kb before the start of the satellite blocks, and reach zero in their vicinity. Note also that AAGAG blocks were barely present in the HiFi assembly (∼0.5 kb), reflecting the much higher bias in these reads. The LILAP assembly fares a little better, but the majority of its assembled AAGAG blocks are very small (median: 80 bp; max: 290 bp; n: 39) and occur at the edges of the contigs (36 out 39 are located at less than 500 bp of an edge); they are the start of large blocks that remained almost completely unsequenced (*e.g.*, Supplemental Fig. S3). It seems clear that these terminal AAGAG blocks disrupted the LILAP contig assemblies due to a lack of reads. Finally, R6 has a very poor representation of all satellites. The second satellite, AAGAGAG is very similar to the first, except for a smaller abundance. The last satellite, AAGGAG, contrasts sharply with the former ones. It is less abundant in the genome and yet is the most abundant in the HiFi and LILAP assemblies (Table 2). More importantly, an 18 kb AAGGAG block has HiFi coverage throughout its length (Supplemental Fig. S4), indicating little or no sequencing bias. Note that the first two satellites are predicted to form perfect (*i.e.*, strong) triplex DNA structures and are very toxic, whereas the last would form a weak triplex DNA, and is benign (Table 2, columns 8-9). This association between triplex-forming potential and sequencing bias – across all three AG-only satellites for which we have reliable coverage data – indicates that triplex-forming, and not GA-richness, is the cause of sequencing bias in AG-only satellites. It is worth noting that AAAGG, the assembly-breaker satellite reported by (Nurk et al. 2022) in the human genome, is also predicted to form a perfect triplex.

**Table 2.** AG-only satellites in the Drosophila genome. Sequence sizes are in kb.

| Satellite | Males haploid | assembled ONT | assembled HiFi | assembled LILAP | assembled R6 | Mirror repeat <sup>c</sup> | Triplex-forming <sup>d</sup> | Triplex score <sup>e</sup> |
| --- | --- | --- | --- | --- | --- | --- | --- | --- |
| AAGAG | 10970 <sup>a</sup> | 178.4 | 0.5 | 2.8 | 5.1 | yes | yes | 60 |
| AAGAGAG | 2975 <sup>a</sup> | 7.0 | 0 | 0.5 | 0.5 | yes | yes | 60 |
| AAGGAG | 658 <sup>b</sup> | 36.9 | 11.7 | 3.9 | 0.3 | no | no | 17 |

<sup>a</sup> Estimated as $\frac{1}{2} X + \frac{1}{2} Y + \text{autosomes}$ (Table 3 in (Lohe et al. 1993)), to make it directly comparable to male raw reads
<sup>b</sup> These satellites were not studied by (Lohe et al. 1993), and were estimated from the ratio in short reads (e.g., AAGGAG / AAGAG) multiplied by the AAGAG amount reported by (Lohe et al. 1993).
<sup>c</sup> Mirror repeat motif, as evaluated by the *nBMST* program (Cer et al. 2013).
<sup>d</sup> Potential to form triplex DNA, as evaluated by the *gquad* program (Ajoge 2015).
<sup>e</sup> Potential to form triplex DNA, as evaluated by the *triplex* program (Lexa et al. 2011). Ranges from 0 to 60, with 60 meaning perfect triplexes.

One interesting way to quantify the effect of satellites is to measure how far HiFi reads penetrate into the satellite blocks (Supplemental Fig. S5; Supplemental Methods). As shown in Fig. 3A (and detailed in Supplemental Table S5), HiFi read coverage typically drops to zero before entering blocks of the toxic satellites AAGAG and AAGAGAG, whereas it penetrated ∼18 kb into the block of the benign satellite AAGGAG.

**Figure 3.**
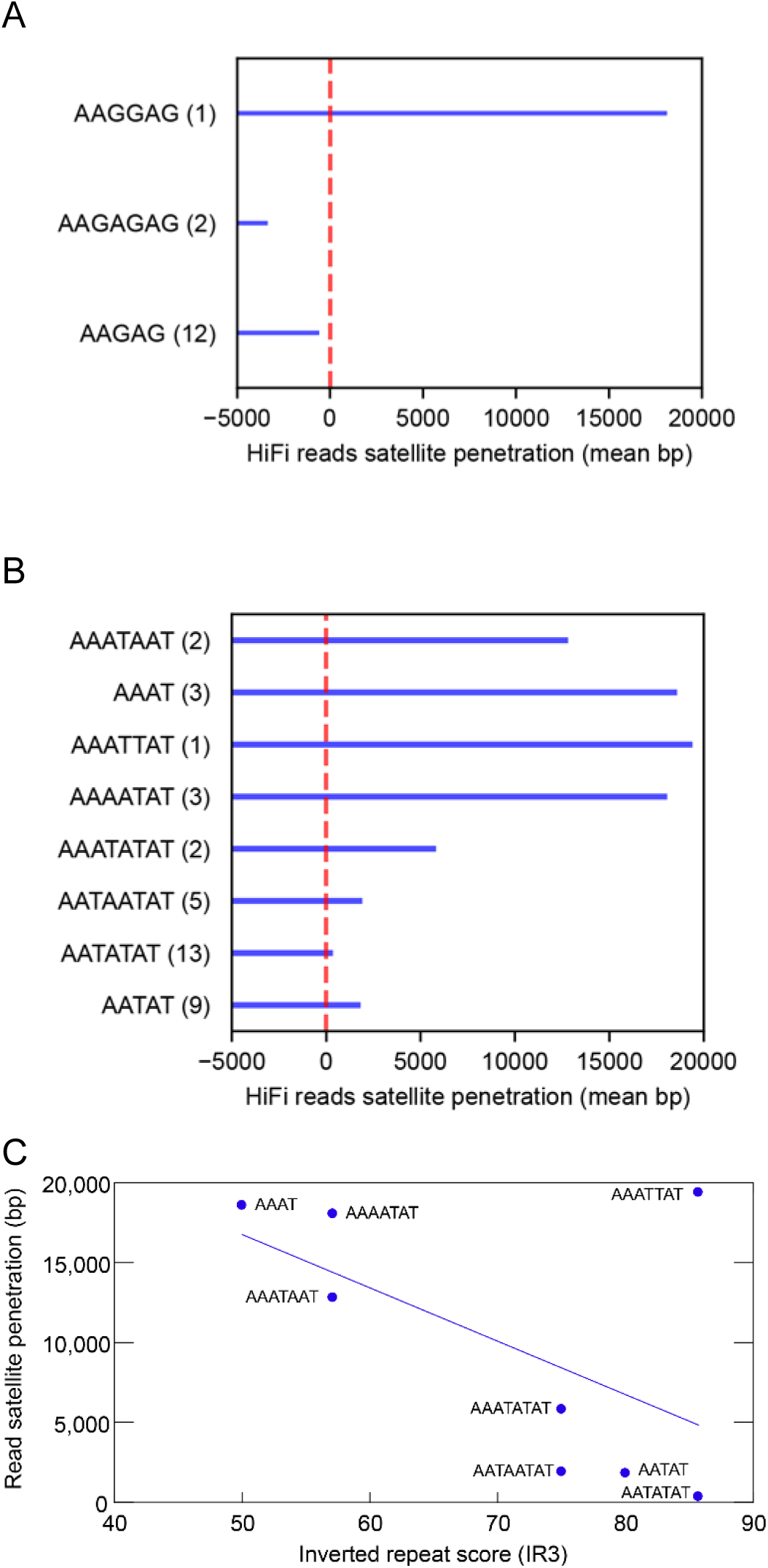
Penetration of HiFi reads into the satellite blocks. A) AG-only satellites. The red dashed line marks the start of the satellite block, and the numbers after the satellite names are the number of blocks used to calculate the average penetration. B) AT-only satellites. C) Relationship between the inverted repeat score and satellite penetration for AT-only satellites. The blue line is the linear regression, which is not statistically significant if we include the outlier AAATTAT (*P* < 10^-3^ excluding it; see text). See Supplemental Table S5 for the raw data.

### How (AATATAT)n and other non-triplex forming satellites cause sequencing bias?

Eight AT-only satellites yielded reliable coverage profiles (Table 3; representative coverage profiles are shown in Supplemental Fig. S6). Four are clearly toxic (AATAT, AATATAT, AATAATAT, and AAATATAT), and four are benign (AAAATAT, AAATTAT, AAAT, AAATAAT). Toxic AT-only satellites are less disruptive than their AG-only counterparts: HiFi read coverage drops to zero ∼2 kb inside the blocks (instead of before entering them; Supplemental Table S5), and significant amounts were assembled using HiFi reads (Table 3). Furthermore, blocks shorter than ∼1kb, or with greater heterogeneity, cause partial but not complete coverage loss (Supplemental Fig. S2).

**Table 3.**
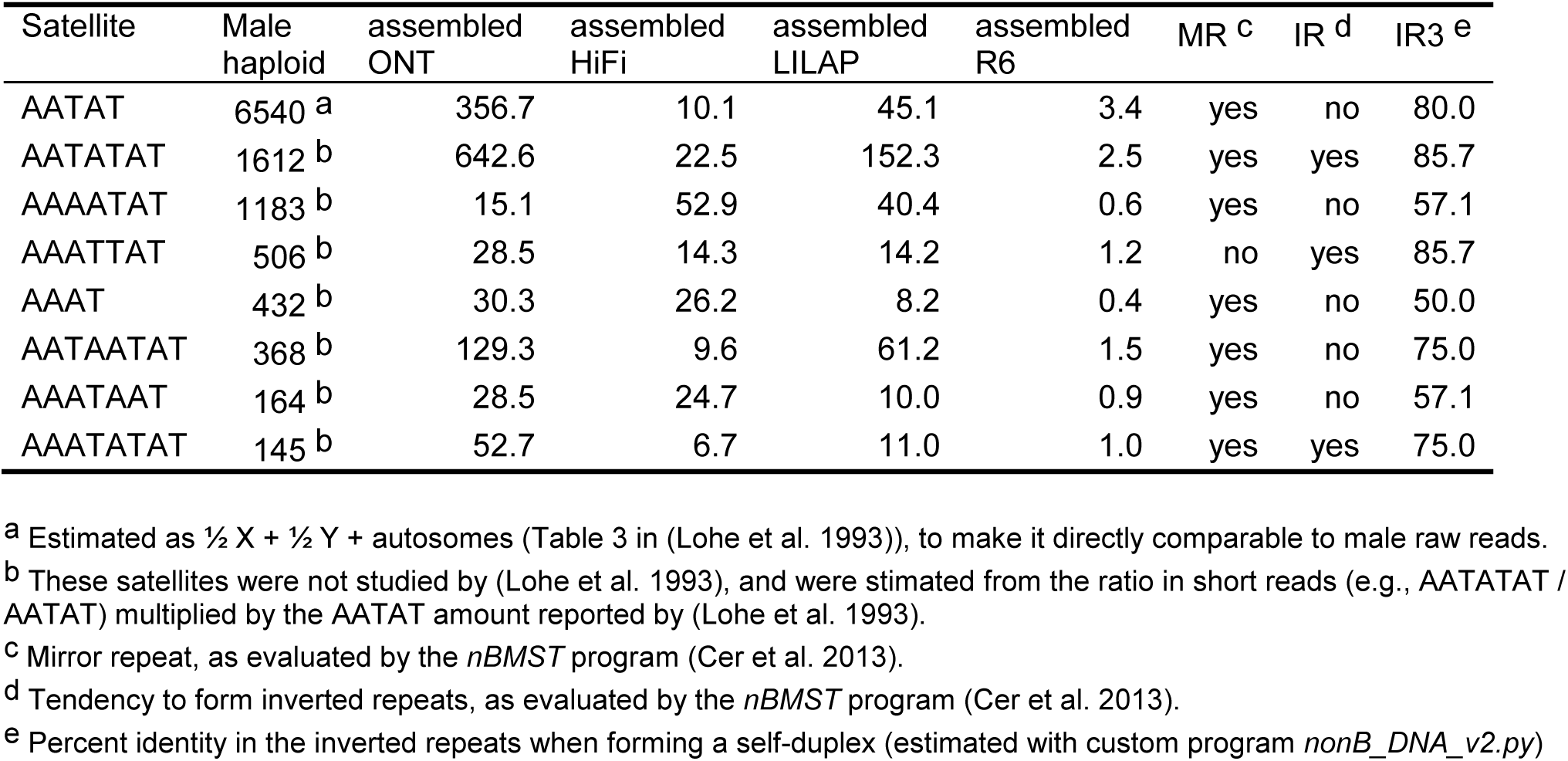
AT-only satellites in the Drosophila genome Sequence sizes are in kb.

Now focusing on AT-only satellites, what explains the difference between toxic and benign ones? Nearly all AT-only satellites are mirror repeats (as are the toxic AG-only satellites AAGAG and AAGAGAG), but none of them can form triplex DNA, because triplex formation additionally requires homopurine / homopyrimidine strands (Mirkin et al. 1987). Mirror repeats *per se* are considered harmless, since formation of triplex DNA is their only known functional property (McGinty et al. 2025). Using the *nBMST* program (Cer et al. 2013) we found that these eight AT-only satellites are also unable to form Z-DNA or G-quadruplex, and that some of them can form inverted repeats (Table 3, column 8). However, the association between inverted repeats (as detected by *nBMST*) and satellite toxicity is poor: only two of the four toxic satellites were flagged as inverted repeats.

Inspired by the *triplex* program (Lexa et al. 2011), we sought to make a quantitative estimation of the strength of the inverted repeats, instead of the usual yes/no output. For this purpose, we calculated a simple score (“IR3”) which ranges from 0% to 100%, with 100% corresponding to a perfect inverted repeat (Supplemental Methods). The IR3 score is shown in column 9 of Table 3, and indicates that the association between inverted repeats and satellite toxicity is quite strong: all four toxic satellites have an IR3 score of 75% or higher, whereas three of the four benign satellites have IR3 <60%. We propose that in the toxic AT-only satellites, the single-stranded DNA produced during ONT or PacBio sequencing folds into long imperfect hairpins (Supplemental Fig. S7; these long hairpins are also called “foldback DNA”; Cech and Hearst 1975). Similarly to the case of triplex DNA and AG-only satellites, these hairpins would form in the vicinity of the DNA polymerase (PacBio) or protein pore (ONT), disrupting sequencing. Note that satellites with an IR3 score of 75% are toxic, implying that long hairpins of AT-only sequences with one mismatch every four bases are stable; there is indeed good support for this (SantaLucia 1998; Peyret et al. 1999; Supplemental Results).

The only apparent exception to the association between satellite toxicity and inverted repeats is the AAATTAT satellite, which behaves as a benign satellite but has a high IR3 score (85.7%). This satellite is represented by a single block close to the *Mitf* gene. It probably should have been excluded from the analysis because this block has signs of misassembly (Supplemental Fig. S8) and is intermingled with other sequences: as shown in Supplemental Table S6 and Supplemental Fig. S9, among satellites with high IR3, the AAATTAT block has by far the smallest uninterrupted length, the lowest proportion of the main satellite, and the highest proportion of other satellites. Hence, it seems that the available data do not allow a reliable classification of its effect on sequencing. We retained this satellite in the analysis to avoid selection bias, by excluding potential counter-evidence. The AAATTAT satellite may also point to aspects of AT-only satellite toxicity unrelated to inverted repeats. For example, a possibly relevant observation is that AAATTAT is the only AT satellite that is not a mirror repeat (Table 3). Although the only known property of mirror repeats is to form triplex DNA (McGinty et al. 2025), which AT-only satellites are unable to do, they may somehow help stabilize imperfect hairpins.

We further explored the relationship between satellite toxicity and inverted repeats by measuring how far HiFi reads penetrate into different satellite blocks. As shown in Fig. 3B (and detailed in Supplemental Table S5 and Supplemental Results), the difference between toxic and benign AT-only satellites is clear-cut. These data allow a statistical test of the association between the IR3 score and satellite toxicity (*i.e.*, penetration by HiFi reads). Including all eight satellites, the association is non-significant (*P* = 0.09; linear mixed-effects regression model), but AAATTAT is a clear outlier (Studentized Residual: 7.5; *P*=0.005; Supplemental Results; Fig. 3C). Excluding AAATTAT, the association between IR3 and satellite toxicity becomes highly significant (*P* < 10^-3^). Exactly the same results were obtained if we replaced IR3 by the predicted thermodynamic stability of the hairpins (the melting temperature *Tm*; Supplemental Results; Supplemental Fig. S10).

Although we classified the satellites in two categories (benign vs. toxic), there are some hints of a continuum. For example, among the benign AT-only satellites, the read penetration of AAAT (IR3 50%) is slightly higher than that of the other benign satellites, AAAATAT and AAATAAT (IR3 57%), suggesting that the latter two are slightly toxic. The melting temperatures of these three satellites are 27°C, 34°C and 34°C (Supplemental Table S7), so it is possible that the last two form hairpins under the conditions used in long-read sequencing. Another hint of this “minor toxicity” is the coverage drop caused by a large block of the benign satellite AAATAAT shown in Fig 1C (IR3 57%; *Tm* 34°C), which seems too strong to be caused only by read mapping issues. These signs of a continuum are hardly surprising, given that the underlying cause of toxicity (the stability of the hairpins) is a continuous variable. The same is probably true for AG-only satellites, as indicated by their heterogeneous effects in the human data (next section).

A final point is that we found that some regions contain satellite blocks with opposite orientations. One of the clearest examples occurred in contig ptg000099l, which contains a 1281 bp perfect AATATAT repeat, an 87 bp spacer, and then 2086 bp of the same satellite in the opposite orientation. The relevance of this phenomenon, if any, remains to be determined.

### Simple satellites in the human genome cause the same type of bias observed in *Drosophila*

Nurk et al. (2022) noted a strong reduction of PacBio HiFi coverage across GA-rich sequences found at a few sites in the human genome. Here we systematically examined this question, by applying the same approach we used for *Drosophila* to the human T2T genome assembled by Nurk et al. (2022). Note that we are looking for perfect repeats; the human genome is known to contain large satellite blocks such as the 27.6 Mbp Hsat3 block in Chromosome 9 (monomer: CATTC; Altemose 2022), but as Carvalho et al. (2026) noted, the longest perfect tandem repeat of CATTC within this block is only 100 bp long. Similar observations hold for other simple human satellites that occur in large blocks. Such highly degenerated satellites affected the HiFi and ONT coverages, but did not cause the major drops that led to assembly breaks (Supplemental Fig. S22 in Nurk et al. 2022).

We found that the human genome has far fewer perfect simple satellites than *D. melanogaster*: there are only 560 individual blocks with homogeneous runs longer than 250 bp, belonging to 425 different monomers, and totaling 250 kb (Table 4). The largest blocks are from the telomeric repeat TTAGGG and are a few kb long; if we remove them, the total length drops to 188 kb. The human Y chromosome contains only 13 simple satellite blocks, with a total length of 12 kb. Across the whole genome, the toxic satellites amount to ∼13 kb, the longest block having 783 bp. These numbers are dwarfed by the *Drosophila* data (Table 4), which is even more striking when we take into account that the human genome is ∼15 times larger.

**Table 4.** Perfect simple satellites in the human and Drosophila genomes. Unless otherwise noted, values are in base pairs.

| Satellite feature | Human | *Drosophila*<sup>b</sup> |
| --- | --- | --- |
| Number of blocks ≥ 250 bp | 560 | 2,648 |
| Number of satellite monomers | 425 | 65 |
| Total length of satellites | 249,631 | 2,662,547 [34 Mb] |
| Longest block of satellites | 3,228 | 56,484 |
| Total length of toxic satellites<sup>a</sup> | 12,833 | 1,366,676 [20 Mb] |
| Longest block of toxic satellites | 783 | 18,008 |

<sup>a</sup> Human toxic satellites: all satellites listed in Supplemental Table S9 and that reduced HiFi average coverage to less than 29 × ; *Drosophila*: AAGAG, AAGAGAG, AATAT, AATATAT, AATAATAT, AAATATAT.
<sup>b</sup> Numbers in brackets are direct measurements taken from (Lohe et al. 1993), adjusted to haploid male genome .

Despite their small size, human satellites do cause coverage drops and assembly breaks (Nurk et al. 2022). In order to study them, we measured the read coverage across all 560 simple satellite blocks (Supplemental Methods; Supplemental Table S8). We found that many telomeric satellite blocks (TTAGGG; represented as AACCCT in our tables) have low coverage, but this is due to the expected reduction in coverage toward the ends of the chromosomes (Supplemental Fig. S11, panel J). Excluding the telomeric satellite blocks, among the ten lowest coverage individual blocks, nine belong to satellites predicted to form perfect triplexes (AAAGG, AAG, AGAGGG, and AGAGGGG; Supplemental Table S8). One of these blocks, a series of closely spaced AAAGG blocks located on chromosome 2 and totaling 1.1 kb, caused a 955 bp zero coverage stretch (coordinates: 67916197-67917151; Supplemental Fig. S11A).

We selected for detailed inspection the 22 satellites with average coverage below half of the genome-wide coverage (29×). We examined their individual block coverage profiles looking for the association between the satellite and coverage drops and also for co-occurring satellites (Fig. 4; Supplemental Fig. S11); blocks where the target satellite occurred intermingled with other satellites were deemed unreliable and excluded from the analysis (Supplemental Table S9). Among these 22 satellites, one is the telomeric repeat TTAGGG, 13 co-occur with other satellites (hence their coverage profiles were unreliable; many form imperfect hairpins), and eight yield reliable coverage profiles with clear coverage drops (Fig. 4; Supplemental Table S9; Supplemental Fig. S11). Four of these eight satellites are predicted to form perfect triplexes (score 60; AAAGG, AAG, AGAGGG, and AGAGGGG), two are predicted to form imperfect hairpins, and two are predicted to form weak and probably irrelevant triplexes (AAGAAGTGGGAGGGGGAGG, triplex score 27; AAGGGAGGAAGGGCAGGGTGGGAGG, triplex score 19). These last two have many G residues in tandem and are predicted to form G-quadruplex DNA (Supplemental Table S9). Hence, it is likely that, in addition to triplexes and hairpins, G-quadruplex satellites can cause HiFi coverage drops.

**Figure 4.**
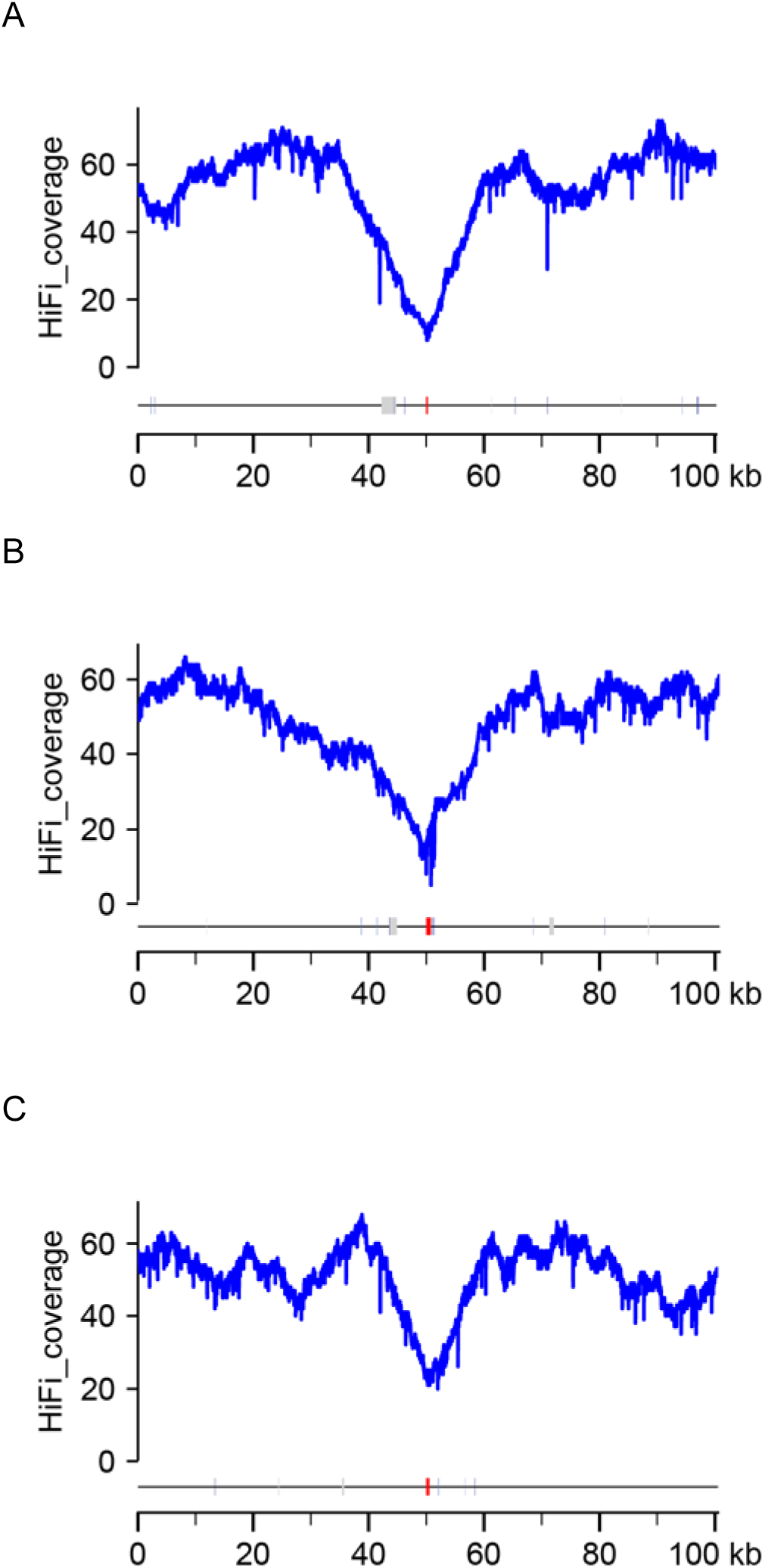
Three simple satellites that cause strong HiFi coverage drops in the human genome. The target satellites are shown in red and transposable elements in light grey. A) a 256 bp block of AAAGG in chromosome 8, predicted to form a perfect triplex. B) a 783 bp block of AAAATGTATATTATATATATTATATAT in chromosome 2, predicted to form an imperfect hairpin (82% identity). C) a 513 bp block of AAGAAGTGGGAGGGGGAGG in chromosome 12. The only non-B DNA structure predicted for this satellite is G-quadruplex.

For comparison with *Drosophila*, we also did a detailed inspection of all human AT-only and AG-only satellites, regardless of their effect on coverage (Supplemental Table S9). As other human simple satellites, AT-only blocks are very small (the largest has 546 bp); almost all are predicted to form imperfect hairpins but are not associated with clear coverage drops. This lack of effect on coverage is not surprising, since the *Drosophila* data show that AT-only satellites require a few kb to confer toxicity (Supplemental Fig. S2). Only one AT-only satellite caused a distinguishable (albeit small) coverage drop: AAATATATATAATATATATTTATAT (Supplemental Fig. S11, panel Y), which is predicted to form a nearly perfect hairpin (IR3: 96%). The nearly perfect hairpin probably explains its toxicity despite the small block size (300 bp).

Regarding the AG-only satellites, we found that perfect triplex satellites have heterogeneous effects in the human data, some of them causing strong drops in coverage (*e.g.*, AAAGG, AAG), and others producing smaller drops (*e.g.*, AAGGG) or no clear drop at all (*e.g.*, AAAAG; Supplemental Fig. S11, panels A, B, D, Q, W, X, AC). This heterogeneity is most likely explained by the small size of the blocks and the fact that the triplex score is a proxy for the relevant parameter, namely the thermodynamic stability of the triplexes. Unfortunately, as might be expected from three DNA strands associating via Watson-Crick and Hoogsteen base pairing, the thermodynamics of triplex formation is complex and not completely understood, being affected by size, GC-content, the geometry of the triplex (parallel vs. anti-parallel), pH, ionic strength, and other factors (Zhang et al. 2025; Santisteban-Veiga et al. 2026). It seems likely that AAAAG and similar satellites would form less stable triplexes that would require more than 405 bp (the size of the largest AAAAG block) to become toxic. Whatever the case, the findings on the human genome are in general agreement with *Drosophila*: strong drops in HiFi coverage are caused by simple satellites that form non-B structures: strong triplexes, hairpins, and perhaps G-quadruplexes (the latter found only in the human data).

### Is read initiation or read failure the immediate cause of sequencing bias?

While investigating the regions of the *Drosophila* genome close to toxic simple satellites, Carvalho et al. (2026; their Table 4) found that ONT reads seldom start within satellite blocks, despite they comprising a large fraction of the sequence. They concluded that satellite DNAs somehow suppress read initiation. The HiFi coverage data reported here on the human genome prompted us to rectify this conclusion. The point is that, given the small size of the blocks of toxic satellites in the human genome, the chance of a DNA fragment starting within them is equally small, so one might expect that only a small number of potential reads would be suppressed. This small number of suppressed reads seems incompatible with the deep and wide drop in HiFi coverage observed in the human data (Fig. 4). An alternative explanation is that DNA fragments containing toxic satellite DNA at any position would cause sequencing failure and thus be absent from the raw read datasets. As a simple test, we simulated HiFi reads of the human genome region shown in Fig. 4A, under these two hypotheses, and obtained their coverage profiles. Indeed, the comparison between the real data (Fig 4A) and simulated coverage profiles (Fig. 5; Supplemental Code) shows that sequencing failure, and not read initiation suppression, is the likely immediate cause of the sequencing bias. Additional confirmation may be obtained by examining the HiFi failed reads (the “*. scraps.bam” file); we would expect to find failed subreads stopping within satellite DNA. It is interesting to note that with long blocks of toxic satellites (as occur in *Drosophila*), the two simulated coverage profiles are more similar (Supplemental Fig. S12). This, and the fact that Carvalho et al. (2026) examined only exon coverage, explain their mistaken conclusion.

**Figure 5.**
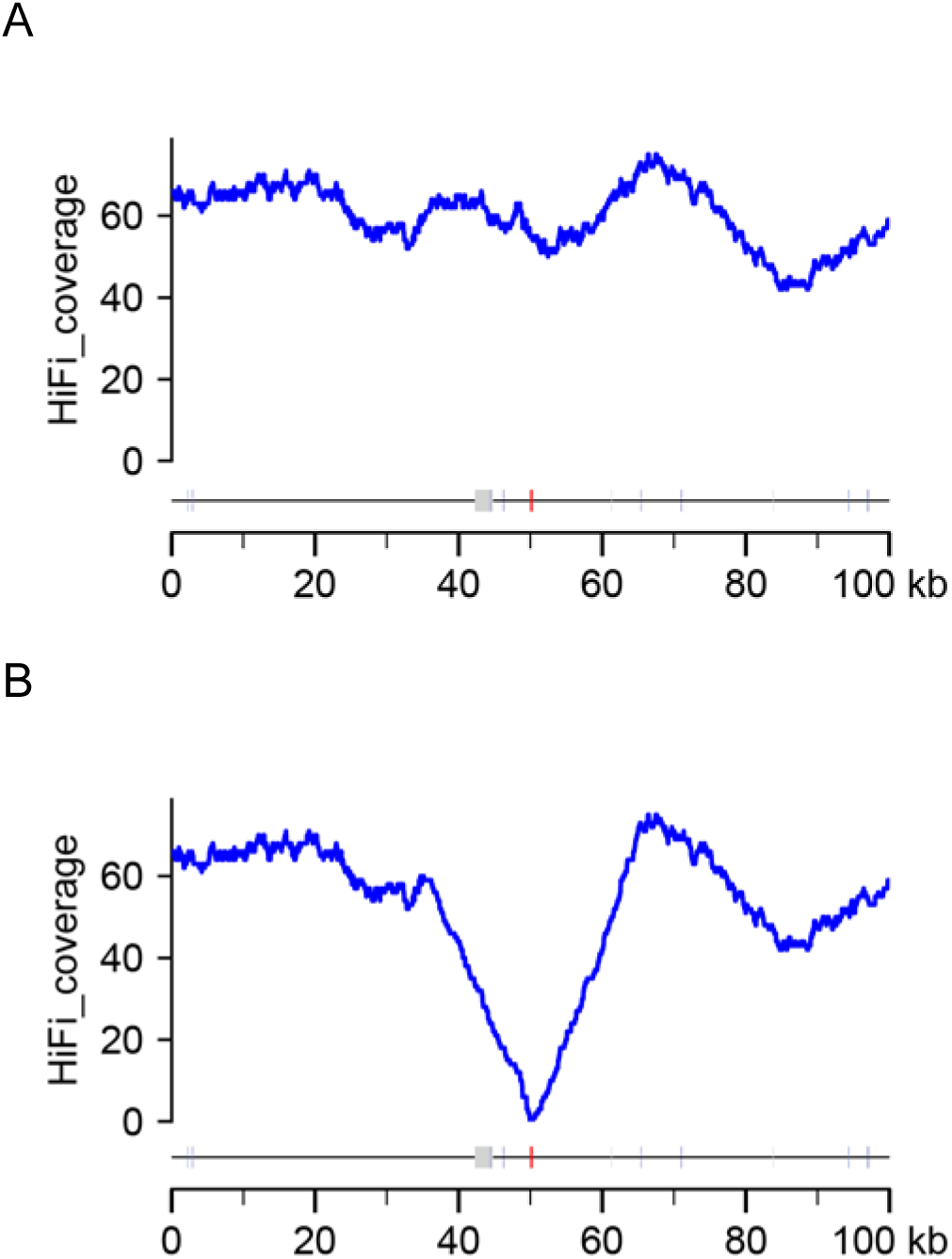
Simulation of sequencing bias: read initiation vs read failure (human genome). We simulated HiFi reads (15 kb reads; 58 x coverage) of the region of human chromosome 8 encompassing the 256 bp AAAGG satellite block (Fig. 4A), then computationally applied two types of bias, and measure read coverage. A) bias caused by reads starting within the AAAGG block being lost. Note that coverage is essentially constant. B) bias caused by reads containing 200 bp or more of AAAGG repeats in any position being lost. Note the deep and wide coverage drop, very similar to the real data shown in Fig. 4A. Transposable elements are shown in light gray and the AAAGG satellite in red.

Strictly speaking the above conclusions apply only to HiFi reads; blocks of toxic satellites in the human genome are too small to affect ONT coverage (Supplemental Fig. S1). But as noted before, the consistency of the bias across PacBio CLR, PacBio HiFi, LILAP, and ONT datasets suggests that it has the same underlying causes across all long-read sequencing platforms (Carvalho et al. 2026).

### Can a single-strand DNA binding protein mitigate the sequencing bias?

If the sequencing bias is caused by non-B DNA structures formed from single-stranded DNA produced during sequencing, then sequestering or digesting this single-stranded DNA should mitigate the bias. We tested this by adding single-strand DNA binding protein (SSB) to ONT sequencing libraries prepared from *D. melanogaster* iso-1 males, at increasing concentrations (Methods). At low concentration (0.5 ug/mL) sequencing proceeded normally, but we observed no difference in raw read satellite content between the SSB-treated library and the matched no-SSB control (Supplemental Fig. S13). At higher concentrations (1 ug/mL and 5 ug/mL) the majority of pores were lost within minutes of loading, and were not recovered even after extensive washing of the flow cell; no usable data were obtained. The addition of bovine serum albumin to the sequencing mix (“BSA blocking”) did not prevent the damage of the flow cells. Thus, SSB has no detectable effect at concentrations the flow cells tolerate and destroy them at concentrations that might have an effect. This side-effect prevented the test of the SSB effect on the satellite bias.

It should be noted that this was a pilot experiment using only ONT, and one single-strand DNA removal approach. It would be most desirable to test SSB in HiFi sequencing, and single strand nucleases in both technologies, but as we argued in the Discussion, this would be better done by the sequencing companies.

### Systematic base-call errors in ONT sequencing

ONT, HiFi, and LILAP assemblies often differ in the extent to which they penetrate satellite blocks. HiFi and LILAP, being more satellite-sensitive, usually stop earlier, and their contigs are typically contained within the longer ONT contig (*e.g.*, Supplemental Fig. S3). We found, however, that LILAP and ONT contigs covering the *CCY* exon 2 region contain different satellite sequences: LILAP has an AAGAC block, whereas ONT has AAAC (HiFi failed to assemble this region). A similar discrepancy occurs near a *kl-2* exon (Supplemental Fig. S14). This might be a simple misassembly, but two lines of evidence suggest otherwise: (i) The HiFi read SRR29479668.730520 spans the whole region in both assemblies, from the *CCY* exon to the start of the satellite block; (ii) The AAAC satellite is completely absent from Illumina reads of the same strain, whereas AAGAC is abundantly present. These evidences strongly indicate that the AAAC satellite in the ONT assembly results from a basecall error of the AAGAC satellite. Basecall errors frequently occur in a strand-specific pattern (Guo et al. 2012; Tan et al. 2022). Consistent with this, the AAAC satellite in ONT reads has a strongly skewed forward / reverse ratio of 41 (1,025,008 bp / 24,780 bp); in contrast, the AAGAC satellite in HiFi and Illumina reads of the same strain shows essentially balanced ratios (respectively, 14,4 Mbp / 17,8 Mbp and 10,49 Mbp / 8,4 Mbp). The AAAC miscalling is most likely analogous to the ONT basecall errors in telomeric repeats reported by (Tan et al. 2022). Carvalho et al. (2026; their Fig. 3) listed AAAC as a common satellite near some Y-linked exons; we now recognize this as a basecall artifact.

How common are miscalled satellites in long-read datasets? The forward / reverse ratio, and presence / absence in Illumina reads provide a “suspects list” that can later be confirmed by aligning HiFi and ONT assemblies (*e.g.*, Supplemental Fig. S14). As shown in Supplemental Fig. S15, Illumina and HiFi reads show a tight correlation in forward-reverse satellite abundance, whereas ONT reads show a wide dispersion. Using a stringent criteria of 10-fold forward / reverse ratio and complete absence in Illumina reads, 11 satellites detected in ONT reads (out of 487) would be the result of base-call errors. When we restricted the analysis to satellites present in the ONT-hifiasm assembly, three of the 65 detected satellites met the criteria for basecall errors: AAAC, AATGG, and CG. ONT reads thus have systematic basecall problems with several simple satellites. These errors could potentially be addressed by tweaking the basecaller, as has been done for telomeric repeats (Tan et al. 2022). In the meantime, although ONT assemblies have clear advantages such as reduced sensitivity to toxic satellites and longer reads, it is safer to cross-check ONT-based results against Illumina and HiFi data.

### Brute force fails to solve the satellite bias problem

Genomes are typically sequenced at 50× to 100× coverage, and one might hope that several-fold deeper coverage would eventually circumvent the sequencing bias and fill the gaps. Note, however, that this bias was discovered in our previous study despite 400× coverage of the *Drosophila* genome (200× in the sex chromosomes; Carvalho et al. 2026). The recent publication of an ultra-high coverage Canton-S assembly (967x Ultra-Long Nanopore, 107x HiFi, plus Hi-C Illumina reads; Liu et al. 2025) allowed for a robust test of the brute force approach. Liu et al. (2025) closed many heterochromatic gaps (18 gaps remained, according to the authors) and described the result as “nT2T” (for “near telomere-to-telomere”). As we showed elsewhere (Carvalho et al. submitted to Nature Communications), this “near completeness” did not hold upon closer inspection: the Canton-S HiFi and ultra-long ONT data suffer from the same bias imparted by simple satellites, and the nT2T assembly actually is missing at least 25 protein-coding genes; overall, it is closer to an incremental improvement than to a T2T assembly. The conclusion is that brute-force will not achieve a complete *Drosophila* genome; the sequencing technologies must be improved.

## Discussion

Sequencing bias against simple satellites is an interesting phenomenon with very undesirable consequences. It caused assembly breaks in the initial human T2T efforts (circumvented by the use of the more robust ONT sequencing; Nurk et al. 2022), and as we argue below, is the main obstacle to a *Drosophila* T2T assembly. It is also bound to appear repeatedly, as the frontier of genomics moves toward T2T assemblies in a wide range of organisms. Overall, long-read sequencing seems to be uniquely susceptible to non-B DNA because it generates long single-stranded DNA in non-denaturing conditions, so structures that would not form in double-stranded DNA (due to competition from regular Watson-Crick pairing) are now energetically favored.

### Sequencing bias against simple satellites is caused by triplex DNA and inverted repeats

In this paper, we showed that the sequencing bias is caused by two distinct non-B DNA structures: triplex-DNA formed by some AG-only satellites, and imperfect hairpins formed by some AT-only and AT-rich satellites. Both affect only the subsets of satellites predicted to form strong non-B structures, providing compelling evidence that non-B DNA is the cause, rather than base composition. All long-read technologies are affected, PacBio HiFi reads being the most sensitive, followed by PacBio LILAP, and then ONT (Carvalho et al. 2026; Supplemental Fig. S1). On the other hand, ONT is uniquely prone to miscalling some simple satellites (Supplemental Fig. S14; Supplemental Fig. S15; Tan et al. 2022), a problem that must be addressed: besides introducing systematic errors in an assembly, it will most likely fragment it, by disrupting the detection of read overlaps.

Triplex-forming AG-only satellites cause a powerful sequencing bias: HiFi coverage usually drops to zero before the start of long satellite blocks (Fig. 2A), and a block as small as 256 bp of AAAGG is enough to cause near zero coverage (Nurk et al. 2022; Fig. 4). AT-only satellites are less toxic than their AG-only counterparts: HiFi coverage usually only drops to zero inside the blocks (Fig. 2B), and a 1 kb block of the toxic satellite AAATATAT only reduces the coverage to half (Supplemental Fig. S2). Why would AT-only satellites be less toxic than AG-only ones? It is possible that hairpins are intrinsically less disruptive for sequencing than triplex DNA. An alternative explanation is that all AT-only satellites we found form imperfect hairpins, whereas toxic AG-only satellites form perfect triplexes. The presumably weaker imperfect hairpins would make the toxicity of AT-only satellites more fickle, requiring longer, more homogeneous blocks for full effect. Many satellite blocks we found have some degree of heterogeneity (deletions, insertion of non-satellite sequences, *etc.*). Given the larger size requirement of AT-only satellites, this heterogeneity is expected to disproportionately attenuate their toxicity, when compared to AG-only satellites.

Irrespective of their effects on sequencing, we may ask why all inverted repeat AT-only satellites are imperfect. A possible explanation (besides chance) is that an Mbp-sized perfect inverted repeat could adopt a cruciform DNA configuration *in vivo* and would therefore not be tolerated. For example, in *Saccharomyces cerevisiae* a 70 bp perfect inverted repeat causes frequent double-strand breaks during meiosis (Nasar et al. 2000). Triplex DNA formation, on the other hand, requires single-strand DNA or unwinding of the double-helix, which only occurs *in vivo* at specific processes (*e.g.*, during DNA replication), and usually in the presence of DNA single-strand binding proteins.

### Simple repeats in the *Drosophila* and human genomes, and the prospect of a *Drosophila* T2T assembly

In general, the findings on the human genome corroborate the *Drosophila* results: all satellites that cause HiFi coverage drops are predicted to form strong non-B DNA structures (triplexes, hairpins, and possibly G-quadruplexes). There is, however, a major difference between the two genomes in the amount of simple satellite DNA, which has been known for a long time. For example, Miklos and John (1979) commented that 4% of the human DNA is composed of satellites, compared with nearly 30% of the *Drosophila* genome. However, when we looked at perfect simple satellites, which is the class able to disrupt long-read sequencing, the difference became staggering: the amount of perfect simple repeats in the *D. melanogaster* genome is several orders of magnitude larger than the human genome (Table 4). The difference shown in Table 4 is even more striking when we take into account that the human genome is ∼15 times larger and is represented by a T2T assembly, whereas the available *Drosophila* genomes are draft assemblies that severely underrepresent simple satellites due to the sequencing bias and assembly collapse. (Lohe et al. 1993) estimated that AAGAG, AAGAGAG, and AATAT - three satellites that disrupt long-read sequencing - account for 20.5 Mbp of the *Drosophila* male haploid genome. These sequences are very abundant in the Y but occur on all chromosomes (Lohe et al. 1993). The human genome, on the other hand, contains ∼13 kb of toxic satellites (Table 4). In many cases a single intron of one *Drosophila* Y-linked gene contains more sequencing disrupting satellites than the whole human genome (*e.g.*, Supplemental Fig. S1; Supplemental Fig. S3). As Carvalho et al. (2026) noted, the finding that several of these very abundant satellites strongly disrupt long-read sequencing uncovers a major hurdle for the achievement of a *Drosophila* T2T assembly. It is no wonder that a *Drosophila* T2T assembly is lagging so much behind the human ones: in a sense, the human genome is an easier target. A *Drosophila* T2T assembly will require correctly assembling across Mbp-sized blocks of satellite DNA, most of them being able to disrupt all current long-read sequencing technologies due to the formation of non-B DNA structures. This is not an impossible task, but as the *Drosophila* Canton-S ultra-deep sequencing shows, brute force will not accomplish it. It certainly will require improvements in the sequencing technologies in order to attenuate or solve the sequencing bias against some simple satellites (as we attempted with as our pilot experiment with single-strand DNA binding protein) and in the case of ONT, to address the satellite miscalling (Supplemental Fig. S14 and Supplemental Fig. S15).

## Concluding remarks

There is now good evidence that the sequencing bias against simple satellites is caused by non-B DNA structures - hairpins and triple helixes - originating from single-stranded DNA. Further computational investigations could clarify some remaining issues (*e.g.*, the possible role of G-quadruplexes), but it seems to us that a direct attack on the bias, by searching for well-tolerated single-strand nucleases or single-strand binding proteins, would be the best approach at this point. This should be done by the PacBio and Nanopore companies because: (i) reagents are all proprietary, so the concentrations of salts, pH, *etc.* are unknown; (ii) flow cells and SMRT cells are prohibitively expensive for a lab to be experimenting with, using conditions that may kill them. Although we reported here a preliminary and unsuccessful attempt at doing this, developing effective solutions is ultimately within the remit of the sequencing companies themselves. We hope they fulfill it, so further advances in *Drosophila* and general genomics become possible.

## Methods

### Raw reads

The raw reads used in this work are listed in Supplemental Table S10. Unless otherwise noted, all data sets were obtained from adult *D. melanogaster* males of the reference strain iso-1.

### SSB sequencing experiments

The experiments were carried out independently in two laboratories (UFRJ/Brazil and Princeton/USA) in Nanopore 10.4.1 flow cells. ONT libraries were prepared from iso-1 males using the SQK-LSK114 protocol. Recombinant *E. coli* single-stranded DNA binding protein (SSB; Promega M3011) was serially diluted in 10 mM Tris-HCl and added to the loading mix immediately before flow cell loading. Final SSB concentrations of 0.5, 1, and 5 µg/mL were tested. BSA blocking was tested with a a SSB concentration of 1 ug/mL; 5 µL of 50 mg/mL BSA was added to the 1170 µL FB / 30 µL FLT priming mix prior to priming, following the manufacturer’s recommendations.

### Genome assemblies

PacBio HiFi and PacBio LILAP reads were assembled using *hifiasm* (Cheng et al. 2026) with default parameters. Nanopore 10.4 reads were assembled using *hifiasm* (setting --ont) and a procedure to increase the representation of the Y chromosome that will be detailed elsewhere. Briefly, we started with the raw reads from Kim et al. (2024), which have ∼400x coverage, and removed the adaptors with Porechop_ABI (Bonenfant et al. 2023). Most genome assemblers work well around 100× coverage, and very high coverages can be detrimental, so we downsampled the reads taking the 100× larger ones (we discarded reads smaller than 45 kb). However, Y-linked reads from biased regions are smaller on average (Fig. 4 in Carvalho et al. 2026), so the above procedure would inadvertently reduce representation of the most biased regions of the *Drosophila* genome. We circumvented this by running the YGS program (Carvalho and Clark 2013) to identify putative Y-linked reads among the reads below 45 kb, and adding those back to the previously size-selected read set. We then run *hifiasm* with the parameters --primary -l 1 --ont. The ONT-*hifiasm* assembly has a much better representation of the satellite blocks and was used in most analyses (Supplemental Fig. S3). For one benign satellite (AAAATAT), the ONT assembly showed signs of repeat collapse in one contig, and was replaced by the HiFi assembly. All genome assemblies are available at Zenodo (https://doi.org/10.5281/zenodo.21311635).

### Detection of repetitive sequences

We ran *Censor* (Kohany et al. 2006) locally to detect transposable elements. As in Carvalho et al. (2026), simple satellites were detected with the custom Python script *find_tandem_repeats_v2.py*, which is based on a regular expression that finds any perfect tandem repeat (head-to-tail) with up to 30 bp monomer size that is present in a DNA sequence (McGinty et al. 2025). For genomic sequences we used a minimum size cut-off of 250 bp of intra-contig perfect repeats or 30 bp at the edges of a contig. We adopted the 250 bp cut-off given that a 256 bp AAAGG sequence caused an assembly break in the human genome (Nurk et al. 2022). The inclusion of terminal 30 bp blocks is irrelevant for the ONT assembly, but important for proper analysis of LILAP and HiFi assemblies, where the most toxic satellites tend to occur at the edges of contigs, in blocks that are seldom larger than 200 bp (their actual sizes are probably much larger). When analyzing raw reads, we reduced the cut-off to 50 bp of perfect repeats to account for the size of Illumina reads (150 bp long) and the higher error rate of raw reads.

### Read coverage estimation

Carvalho et al. (2026) analyzed read coverage in CDS sequences using a BLASTN approach. Here we analyzed coverage in satellite-rich intronic and intergenic regions using full genome assemblies as the reference. For this purpose *minimap2* (Li 2018) is the ideal tool. Most figures (*e.g.*, Fig. 1 and Fig. 2) were produced with the wrapper custom script *coverage_anonymous_graph_v1.sh*, which extracted coverage data from bam files previously obtained with *minimap2*, then ran *Censor* and *find_tandem_repeats_v2.py* (to identify transposable elements and simple satellites, respectively), and then passed all the data to the Python script *visualize_repeats_censor5.py*. In the R6 assembly we focused on the annotated gene regions (file dmel-all-gene_extended2000-r6.33.fasta.gz, downloaded from FlyBase), and in the remaining assemblies we used all satellite-containing regions. All scripts are available at GitHub (https://github.com/bernardo1963/missing_exons_2), and the command lines are reproduced in the Supplemental Code file.

### Statistical procedures

The linear regressions used to test the association between inverted repeat scores and read penetration were performed in R (R Core Team. 2021). The command lines are available in the Supplemental Code file. The generation of read penetration data is described in the Supplemental Methods.

### Evaluation of non-B DNA formation potential

Most simple satellite blocks are interrupted by small amounts of other sequences, including other satellites. In order to unambiguously identify the causative sequences of non-B DNA structures, we evaluated pure sequences of the main satellites generated as follows. From a list of target simple satellite monomers, an *awk* script generated a multi-fasta file containing 250 bp of the satellites (*e.g.*, the 5 bp AAGAG monomer would be repeated 50 times). This multi-fasta file was then subjected to the programs that detect non-B sequences: *triplex* (Lexa et al. 2011), *nBMST* for inverted repeats, G-quadruplex, mirror repeats, Z-DNA (Cer et al. 2013), and *gquad* for triplex, G-quadruplex, and Z-DNA (Ajoge 2015). As explained in the main text and detailed in the Supplemental Methods, the “yes/no” output of *nBMST* was insufficient for the evaluation of inverted repeats; we wrote a Python script (*nonB_DNA_v2.py*) that produces a simple quantitative score (“IR3”) for a list of satellite monomers, and is also a wrapper for the *MELTING 5* program (Dumousseau et al. 2012).

## Data access

The Nanopore raw reads generated in this study have been submitted to the NCBI BioProject database (https://www.ncbi.nlm.nih.gov/bioproject/) under accession numbers SRR39225480 and SRR39225481. All the essential computing codes, examples of their usage, related programs, scripts, and data files are available at GitHub (https://github.com/bernardo1963/missing_exons_2), and in the Supplemental Code file. All genome assemblies were deposited at Zenodo (https://doi.org/10.5281/zenodo.21311635).

## Acknowledgements

We thank Paulo Paiva for help with statistical analysis. This research was funded by FAPERJ–Fundação Carlos Chagas Filho de Amparo à Pesquisa do Estado do Rio de Janeiro, grant CNE2018; CNPq–Conselho Nacional de Desenvolvimento Científico e Tecnológico, grant INCT-EM; and Wellcome Trust, grant 207486Z17Z, to A.B.C.. F.U. is supported by CAPES–Coordenação de Aperfeiçoamento de Pessoal de Nível Superior, Finance Code 001.

## Author contributions

Conceptualization and methodology were by A.B.C. Investigation and formal analysis were by A.B.C. Data production was by A.B.C., F.U., and B.Y.K. Data curation was by A.B.C. Writing of the original draft preparation was by A.B.C. Reviewing and editing were by A.B.C., F.U., and B.Y.K. Funding was by A.B.C. and B.Y.K. All authors have read and agreed to the submitted version of the manuscript.

## References

Ajoge HO. 2015. gquad: prediction of G quadruplexes and other non-B DNA motifs. Vol 2.1–2.

Altemose N. 2022. A classical revival: Human satellite DNAs enter the genomics era. Semin Cell Dev Biol 128: 2–14.

Bonenfant Q, Noe L, Touzet H. 2023. Porechop_ABI: discovering unknown adapters in Oxford Nanopore Technology sequencing reads for downstream trimming. Bioinform Adv 3: vbac085.

Carvalho AB, Clark AG. 2013. Efficient identification of Y chromosome sequences in the human and *Drosophila* genomes. Genome Res 23: 1894–1907.

Carvalho AB, Dupim EG, Goldstein G. 2016. Improved assembly of noisy long reads by k-mer validation. Genome Res: 1710–1720.

Carvalho AB, Emerson JJ, Uno F, Chakraborty M, Shukla HG, Larracuente A, Chang C, Karpen G, Bachtrog D, Kim BY et al. The ‘near T2T’ *D. melanogaster* assembly has missing genes and highly incomplete chromosomes. [Manuscript submitted to Nature Communications].

Carvalho AB, Kim BY, Uno F. 2026. Strong bias in long-read sequencing prevents assembly of *Drosophila melanogaster* Y-linked genes. Genome Res 36: 71–82.

Cech TR, Hearst JE. 1975. An electron microscopic study of mouse foldback DNA. Cell 5: 429–446.

Cer RZ, Donohue DE, Mudunuri US, Temiz NA, Loss MA, Starner NJ, Halusa GN, Volfovsky N, Yi M, Luke BT et al. 2013. Non-B DB v2.0: a database of predicted non-B DNA-forming motifs and its associated tools. Nucleic Acids Res 41: D94–D100.

Cheng H, Qu H, McKenzie S, Lawrence KR, Windsor R, Vella M, Park PJ, Li H. 2026. Efficient near-telomere-to-telomere assembly of nanopore simplex reads. Nature doi:10.1038/s41586-026-10105-6.

Dumousseau M, Rodriguez N, Juty N, Le Novere N. 2012. MELTING, a flexible platform to predict the melting temperatures of nucleic acids. BMC Bioinformatics 13: 101.

Guo Y, Li J, Li CI, Long J, Samuels DC, Shyr Y. 2012. The effect of strand bias in Illumina short-read sequencing data. BMC Genomics 13: 666.

Hisey JA, Masnovo C, Mirkin SM. 2024. Triplex H-DNA structure: the long and winding road from the discovery to its role in human disease. NAR Mol Med 1: ugae024.

Hoskins RA, Carlson JW, Wan KH, Park S, Mendez I, Galle SE, Booth BW, Pfeiffer BD, George RA, Svirskas R et al. 2015. The Release 6 reference sequence of the *Drosophila melanogaster* genome. Genome Res 25: 445–458.

Jia H, Tan S, Cai Y, Guo Y, Shen J, Zhang Y, Ma H, Zhang Q, Chen J, Qiao G et al. 2024. Low-input PacBio sequencing generates high-quality individual fly genomes and characterizes mutational processes. Nat Commun 15: 5644.

Kim BY, Gellert HR, Church SH, Suvorov A, Anderson SS, Barmina O, Beskid SG, Comeault AA, Crown KN, Diamond SE et al. 2024. Single-fly genome assemblies fill major phylogenomic gaps across the Drosophilidae Tree of Life. PLoS Biol 22: e3002697.

Kim KE, Peluso P, Babayan P, Yeadon PJ, Yu C, Fisher WW, Chin C-S, Rapicavoli NA, Rank DR, Li J. 2014. Long-read, whole-genome shotgun sequence data for five model organisms. Scientific Data 1: 140045.

Kohany O, Gentles AJ, Hankus L, Jurka J. 2006. Annotation, submission and screening of repetitive elements in Repbase: RepbaseSubmitter and Censor. BMC Bioinformatics 7: 474.

Lexa M, Martinek T, Burgetova I, Kopecek D, Brazdova M. 2011. A dynamic programming algorithm for identification of triplex-forming sequences. Bioinformatics 27: 2510–2517.

Li H. 2018. minimap2: pairwise alignment for nucleotide sequences. Bioinformatics 34: 3094–3100.

Liu YN, Gao JJ, Zhuang XL, Wu DD, Sun YB. 2025. Near complete assembly of *Drosophila melanogaster* Canton S strain genome. Nat Commun 17: 329.

Lohe AR, Hilliker AJ, Roberts PA. 1993. Mapping simple repeated DNA-sequences in heterochromatin of *Drosophila melanogaster*. Genetics 134: 1149–1174.

McGinty R, Lyskova A, Mirkin SM. 2025. The origin of mirror repeats in the human genome. Nucleic Acids Res 53.

Miklos GL, John B. 1979. Heterochromatin and satellite DNA in man: properties and prospects. Am J Hum Genet 31: 264–280.

Mirkin SM, Lyamichev VI, Drushlyak KN, Dobrynin VN, Filippov SA, Frank-Kamenetskii MD. 1987. DNA H form requires a homopurine-homopyrimidine mirror repeat. Nature 330: 495–497.

Nasar F, Jankowski C, Nag DK. 2000. Long palindromic sequences induce double-strand breaks during meiosis in yeast. Mol Cell Biol 20: 3449–3458.

Nurk S, Koren S, Rhie A, Rautiainen M, Bzikadze AV, Mikheenko A, Vollger MR, Altemose N, Uralsky L, Gershman A et al. 2022. The complete sequence of a human genome. Science 376: 44–53.

Oyola SO, Otto TD, Gu Y, Maslen G, Manske M, Campino S, Turner DJ, Macinnis B, Kwiatkowski DP, Swerdlow HP et al. 2012. Optimizing Illumina next-generation sequencing library preparation for extremely AT-biased genomes. BMC Genomics 13: 1.

Ozturk-Colak A, Marygold SJ, Antonazzo G, Attrill H, Goutte-Gattat D, Jenkins VK, Matthews BB, Millburn G, Dos Santos G, Tabone CJ et al. 2024. FlyBase: updates to the *Drosophila* genes and genomes database. Genetics 227.

Peyret N, Seneviratne PA, Allawi HT, SantaLucia J, Jr. 1999. Nearest-neighbor thermodynamics and NMR of DNA sequences with internal A.A, C.C, G.G, and T.T mismatches. Biochemistry 38: 3468–3477.

R Core Team. 2021. R: A language and environment for statistical computing. R Foundation for Statistical Computing.

SantaLucia J, Jr. 1998. A unified view of polymer, dumbbell, and oligonucleotide DNA nearest-neighbor thermodynamics. Proc Natl Acad Sci U S A 95: 1460–1465.

Santisteban-Veiga A, Chandrasegaran S, Dominguez-Arca V, Sabin J, Pukala TL. 2026. Thermodynamic and kinetic considerations in DNA triplex formation revealed by ITC. Eur Biophys J 55: 293–307.

Shukla HG, Chakraborty M, Emerson JJ. 2025. Genetic variation in recalcitrant repetitive regions of the *Drosophila melanogaster* genome. Genome Res 35: 2023–2040.

Tan KT, Slevin MK, Meyerson M, Li H. 2022. Identifying and correcting repeat-calling errors in nanopore sequencing of telomeres. Genome Biol 23: 180.

Wei KH, Lower SE, Caldas IV, Sless TJS, Barbash DA, Clark AG. 2018. Variable rates of simple satellite gains across the *Drosophila* phylogeny. Mol Biol Evol 35: 925–941.

Zhang S, Shen J, Guan Z, Luo T, Chu C, Zhu H, Mergny JL, Cheng M. 2025. Deciphering and predicting thermal and pH stabilities of triplex DNA under multifactorial conditions. Angew Chem Int Ed Engl 64: e202507190.

